# A novel triple-action inhibitor targeting B-cell receptor signaling and BRD4 demonstrates preclinical activity in chronic lymphocytic leukemia

**DOI:** 10.1101/2022.05.11.491547

**Authors:** Audrey L. Smith, Alexandria P. Eiken, Dalia Y. Moore, Lelisse T. Umeta, Lynette M. Smith, Elizabeth R. Lyden, Christopher R. D’Angelo, Avyakta Kallam, Julie M. Vose, Tatiana G. Kutateladze, Dalia El-Gamal

## Abstract

B-cell chronic lymphocytic leukemia (CLL) results from intrinsic genetic defects and complex microenvironment stimuli that fuel CLL cell growth through an array of survival-signaling pathways. Novel small-molecule agents targeting the B-cell receptor pathway and anti-apoptotic proteins alone or in combination have revolutionized the management of CLL, yet combination therapy carries significant toxicity and CLL still remains an incurable disease due to residual disease and relapse. Single-molecule inhibitors that can target multiple disease-driving factors are thus an attractive approach to combat both drug resistance and combination therapy-related toxicities. Using SRX3305, a novel small-molecule BTK/PI3K/BRD4 inhibitor that targets three distinctive facets of CLL biology, we demonstrate that SRX3305 attenuates CLL cell proliferation and promotes apoptosis in a dose-dependent fashion. SRX3305 also inhibits activation-induced proliferation of primary CLL cells in vitro, and effectively blocks microenvironment-mediated survival signals including stromal cell contact. Furthermore, SRX3305 blocks CLL cell migration toward CXCL-12 and CXCL-13, major chemokines involved in CLL cell homing and retention in microenvironment niches. Importantly, SRX3305 maintains its anti-tumor effects in ibrutinib-resistant CLL cells. Collectively, this study establishes the preclinical efficacy of SRX3305 in CLL providing significant rationale for its development as a therapeutic agent for CLL and related disorders.

## 1. Introduction

Chronic lymphocytic leukemia (CLL) is characterized by the expansion of mature B-cells which accumulate in the blood, bone marrow (BM), and secondary lymphoid organs such as the lymph node (LN) and spleen. Hallmark features of CLL include amplified B-cell receptor (BCR) signaling, defective apoptosis, profound tumor microenvironment (TME) dependency, and immune dysfunction [1,2]. Novel therapies targeting BCR signaling (e.g., ibrutinib and idelalisib) and B-cell leukemia/lymphoma-2 (BCL-2) family proteins (e.g., venetoclax) have significantly improved outcomes for both previously untreated CLL patients and those with adverse disease (e.g., 17p deletion and/or TP53 mutation or complex karyotype). Unfortunately, patients continue to progress on these agents by developing acquired resistance through multiple mechanisms [3].

Bruton tyrosine kinase (BTK) inhibitors (e.g., ibrutinib, acalabrutinib, zanubrutinib) disrupt effective BCR signaling which is vital to CLL cell proliferation and supportive TME interactions [4,5]. BTK inhibition elicits high response rates and durable remission in CLL patients, including those with del(17p), but does not fully eliminate the disease [3,6]. Studies have shown that persistent activation of PI3K-AKT-mTOR, NF-κB, and/or MYC signaling contributes to acquired resistance to ibrutinib and residual disease [7-10]. Relapse on ibrutinib is regularly due to BTK mutations that prevent inhibitor binding and often lead to the development of more aggressive diseases such as relapse/refractory CLL or Richter Transformation (RT) [11,12] from CLL into diffuse large B-cell lymphoma (DLBCL-RT) [13]. Other inhibitors targeting phosphatidylinositol-3 kinases (PI3K) or anti-apoptotic pathways through BCL-2 inhibition can be used to further disrupt proliferation/survival signaling and defective apoptosis in CLL [12]. However, these therapies often require combination with other agents including monoclonal antibodies to elicit durable responses with varied toxicity [3,12]. Despite these advances, CLL remains an incurable malignancy, and relapse post-BTK and/or BCL-2 inhibitor therapy is a major clinical challenge in need of novel therapies.

Combination strategies targeting different facets of CLL biology used earlier in disease management may result in lower rates of emergence of drug-resistant clones [14,15]. Epigenetic dysregulation plays a central role in cancer pathogenesis [16-18], making strategies that therapeutically target abnormal epigenetic factors an area of active investigation. For instance, aberrations in epigenetic modifiers (e.g., *BRD4, CREBBP*) have been identified as early driving events in lymphomagenesis and relapse-associated events including chemoresistance and immune escape [19]. Bromodomain-containing 4 (BRD4), a member of the bromodomain and extra-terminal (BET) family of epigenetic reader proteins, binds to acetylated histones to facilitate the recruitment of transcriptional machinery [20]. BRD4 is enriched at nearly all active promoters and most active enhancers in both normal and transformed cells. Cancer cells display exceptionally higher BRD4 binding at super-enhancers of genes that are crucial to maintaining cancer cell identity and promoting oncogenic gene transcription [21,22]. By doing so, BRD4 regulates the expression of genes that govern cell growth and evasion of apoptosis in cancer (e.g., *MYC, BCL6, BCL2*, and *CDK4/6*) [22-25]. Furthermore, histone-independent roles for BET proteins are relevant in leukemia/lymphoma cell survival, as BRD4 interacts with acetylated RELA [26] which augments NF-κB signaling, a central mediator of both external TME triggers and cell-extrinsic aberrations in CLL [27]. Currently available small-molecule BET inhibitors competitively bind to the acetyl-lysine recognition pocket of BET bromodomains displacing BET proteins from active chromatin, which in turn reduces associated gene expression [28,29].

Recently, BRD4 was reported to be overexpressed in primary CLL cells when compared to normal B-cells [30]. In that study, BRD4 was enriched at hallmark genes implicated in CLL disease biology and progression such as BCR pathways associated genes (e.g., *BTK, BLK, SYK, PLCG2*, and PIK3CG), *ZAP70, CXCR4, MIR155, IL4R, IL21R, IKZF1*, and *TCL1A* [30]. BET inhibition has demonstrated preclinical activity in various leukemia and lymphoma models [31], including CLL. Novel BET inhibitors (e.g., PLX51107, OTX015, and GS5829) were reported to reduce CLL cell proliferation, induce cell cycle arrest, and promote cell apoptosis *in vitro* even in the presence of TME protection [30,32]. Moreover, BET inhibition significantly decreased tumor burden and prolonged survival in murine models of aggressive CLL [30]. Despite the remarkable preclinical activity of BET inhibitors in hematological malignancies, studies to identify beneficial combination therapy approaches to enhance efficacy have been reported [31]. As expected, synergism was observed when combining BET inhibitors (e.g., BAY 1238097, GS-5829, OTX015) with inhibitors of BTK (ibrutinib), SYK (entospletinib), PI3K (copanlisib, idelalisib), or BCL2 (venetoclax) in preclinical models of CLL [32-34] and DLBCL-RT [35]. Interestingly, the anti-leukemic activity of BET inhibitor, GS-5829, proved to be synergistic in combination with ibrutinib or idelalisib in primary CLL / nurse-like cell co-cultures reflective of the LN TME in CLL [32].

We recently introduced thienopyranone (TP) scaffold-based chemotypes [36] for the combinatorial inhibition of BTK, PI3K-AKT, and BRD4-MYC in a single compound (i.e., SRX3262 and SRX3305) and demonstrate impressive preclinical activity in mantle cell lymphoma (MCL) [37,38]. BTK/PI3K/BRD4 triple inhibitors present a potential solution to the toxicity that often plagues combination therapeutic strategies that are necessary to overcome CLL progression and drug resistance. Importantly, our studies demonstrated that these novel multi-action inhibitors yield potent anti-tumor effects, while sparing healthy bystander cells and are less toxic to healthy donor B-cells than the combination of single-target drugs required to effectively inhibit the same three targets. In this study, we extend our evaluation of SRX3305 to include preclinical models of CLL and DLBCL, including ibrutinib-resistant CLL, further validating the efficacy of a multi-target single-molecule approach to overcome TME- and drug-induced resistance mechanisms in B-cell malignancies.

## 2. Results

### 2.1 SRX3305 inhibits proliferation and induces apoptosis of malignant B-cell lines

The anti-proliferative properties of SRX3305 or SRX3262 (first-generation BTK/PI3K/BRD4 triple inhibitor) were first evaluated in a panel of B-cell non-Hodgkin lymphoma (B-NHL) cell lines representative of CLL (OSU-CLL, HG-3, MEC-1 and MEC-2) and DLBCL (OCI-LY3 and SU-DHL-6). Key cell line characteristics/genetic aberrations are summarized in ***Supplementary Table S1***. SRX3305 or SRX3262 significantly inhibited CLL cell proliferation in a dose-dependent manner (***Figure 1A – D***). We report an average IC_50_ ∼1.3 µM across the CLL lines evaluated, ranging from 300 nM to 2.85 µM. SRX3305 appeared to be superior to SRX3262 (on average ∼3-fold more potent, P<0.05), hence was used for all further studies. SRX3305 is at least equipotent to common single-target inhibitors (ibrutinib, idelalisib, JQ1 or OTX015) evaluated on this panel of B-NHL cells. Similarly, SRX3305 reduced cell proliferation in DLBCL cells (***Supplementary Figure S1***) with IC_50_ values of ∼290 nM in OCI-LY3 cells (activated B-cell DLBCL subtype) and ∼920 nM in SU-DHL-6 cells (germinal center DLBCL subtype). We next demonstrate that SRX3305 induces apoptosis in HG-3 and OSU-CLL cells in a dose-dependent manner at 24 h (***Figure 1E – F***), and a similar trend was witnessed at 48 h (data not shown). This indicates that SRX3305 not only induces a cytostatic effect, but also a cytotoxic effect in CLL cells.

**Figure 1.**
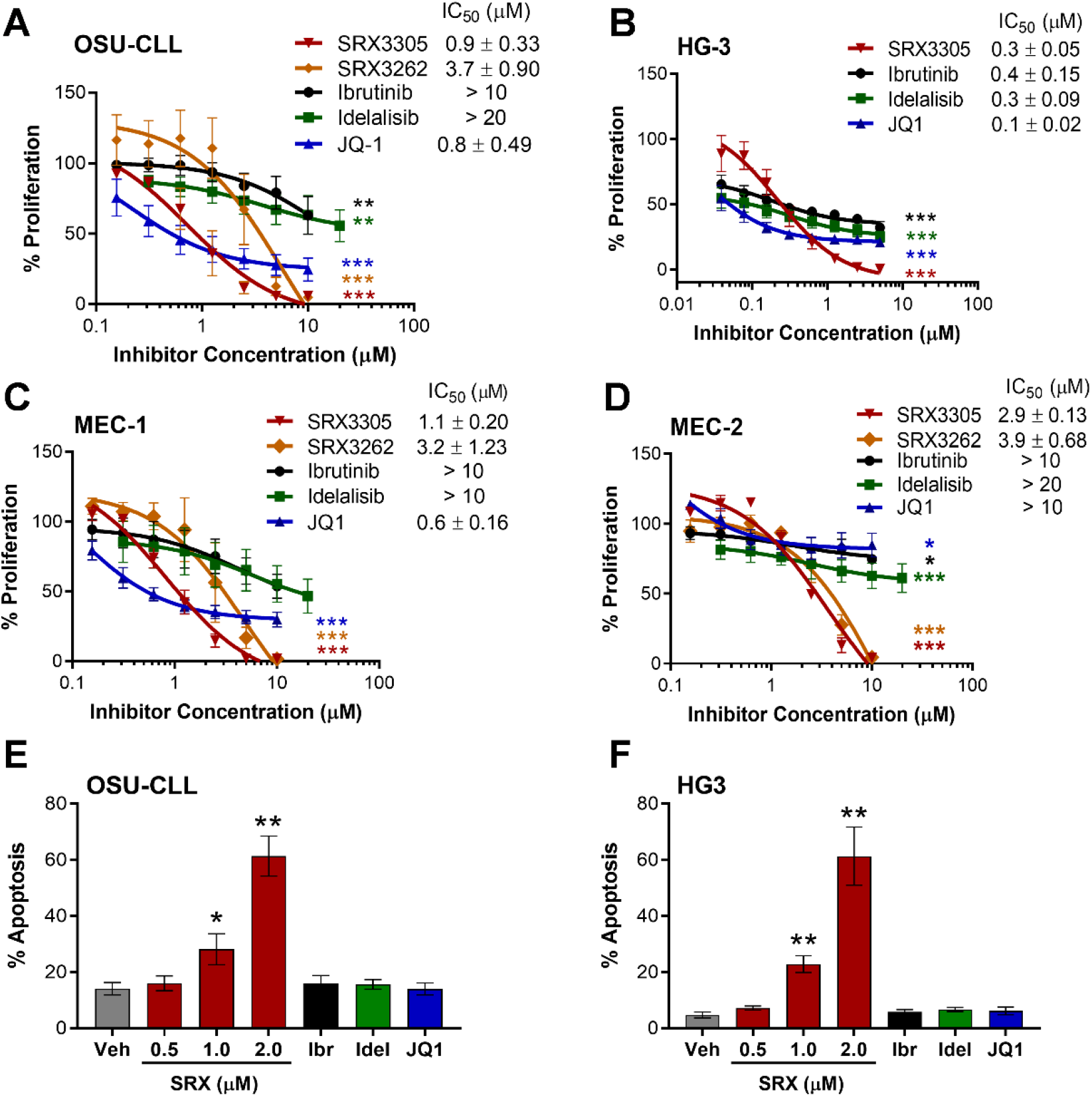
SRX3305 is cytotoxic to CLL cells in a dose-dependent manner. Four CLL cell lines, OSU-CLL **(A)**, HG-3 **(B)**, MEC-1 **(C)**, MEC-2 **(D)**, were treated with increasing concentrations of SRX3262 or SRX3305 (BTK/PI3K/BRD4 triple-inhibitor), ibrutinib (BTK inhibitor), idelalisib (PI3K inhibitor), or JQ1 (BET inhibitor) for 72 h. Proliferation was assessed via MTS assay and results are given as % proliferation normalized to vehicle. Error bars indicate SEM (n=3 independent experiments per cell line). IC_50_ values (mean ± SEM) are noted for each inhibitor within the in-figure legends. Dose-dependent decrease in CLL cell proliferation is observed with asterisks denoting significance *vs*. vehicle. **(E)** OSU-CLL and **(F)** HG-3 cells were treated with DMSO vehicle (Veh), SRX3305 (0.5, 1, 2 µM), ibrutinib (Ibr, 1 µM), idelalisib (Idel, 1 µM), or JQ1 (0.5 µM) for 24 h. Percentage of apoptosis (annexin V-positive cells) was evaluated via flow cytometry (n=3 independent experiments per cell line). Results are given as mean ± SEM. Asterisks denote significant inhibitor-induced apoptosis *vs*. control vehicle. * *P*<0.05, ** *P*< 0.01, *** *P*< 0.001.

### 2.2 SRX3305 inhibits critical BCR survival signaling in CLL

Two major targets of SRX3305, BTK and PI3K, are vital to BCR survival and proliferation signaling [39]. The third, BRD4, is critical for the transcriptional regulation of numerous factors driving CLL pathogenesis, including MYC [23,30]. We show that SRX3305 effectively inhibits the phosphorylation of BTK and PRAS40 (indicative of PI3K/AKT signaling), and reduces MYC expression in OSU-CLL, MEC-1 and MEC-2 cells (***Figure 2A***). Single-target inhibitors (ibrutinib, idelalisib, JQ1) were used as controls.

**Figure 2.**
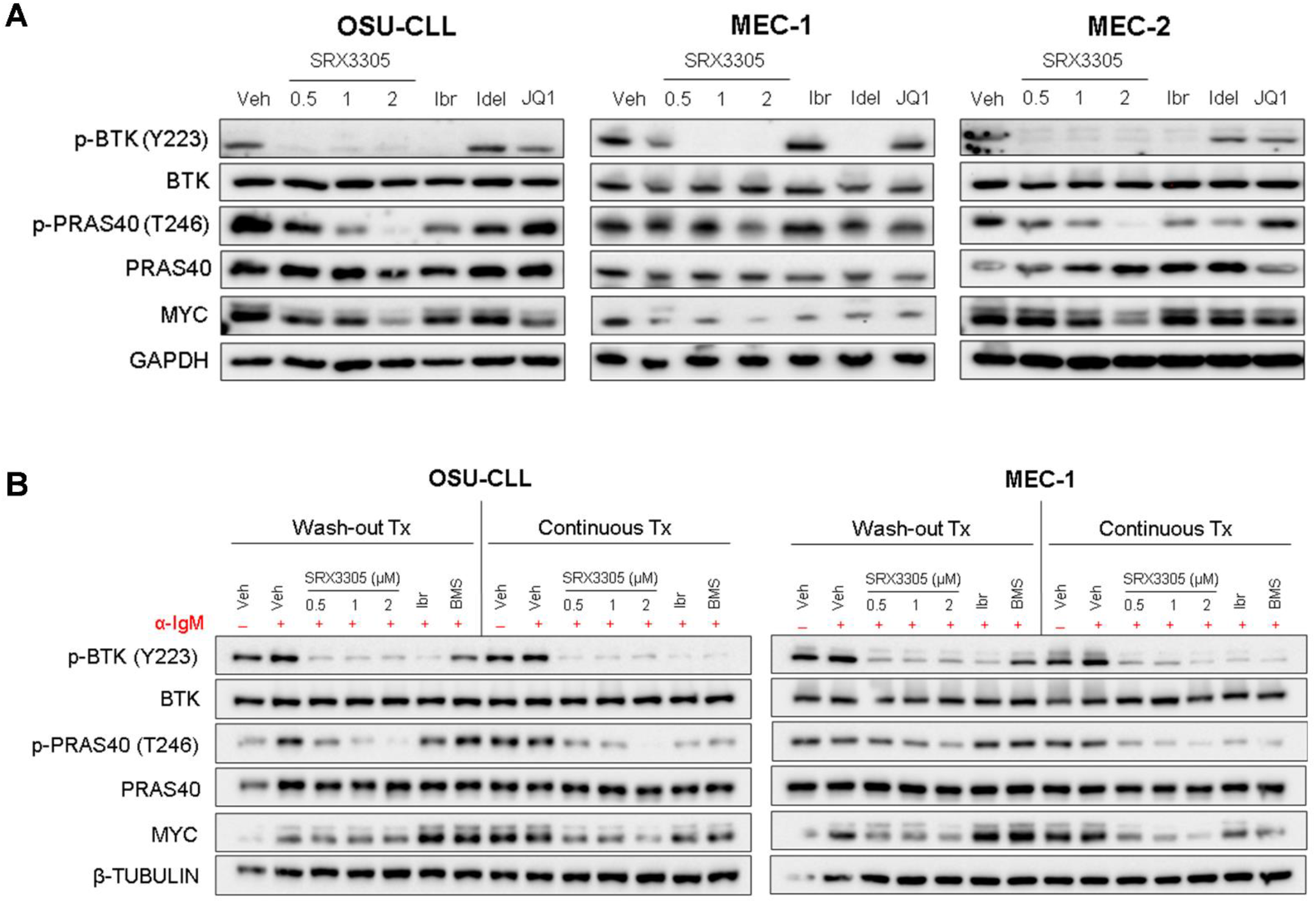
SRX3305 modulates critical B-cell survival pathways in CLL. **(A)** CLL cell lines (OSU-CLL, MEC-1 and MEC-2) were treated with DMSO vehicle (Veh), SRX3305 (0.5, 1, 2 µM), ibrutinib (Ibr, 1 µM), idelalisib (Idel, 1 µM), or JQ1 (0.5 µM). After 4 h, whole-cell lysates were collected and analyzed for phosphorylation and expression of the given proteins, indicative of modulation of BTK (p-BTK/BTK), PI3K (p-PRAS40/PRAS40), and BET/BRD4 (MYC). GAPDH serves as a loading control (n=3 independent experiments per cell line). Representative immunoblots are shown. **(B)** OSU-CLL and MEC-1 cells were treated with DMSO vehicle (Veh), SRX3305 (0.5, 1, 2 µM), 1 µM ibrutinib (Ibr; irreversible BTK inhibitor), or 1 µM BMS-986142 (BMS; reversible BTK inhibitor). Cells were either incubated with treatment continuously for 4 h (Continuous Tx), or treatment was washed out after 1 h and replaced with fresh media for the remaining 3 h of incubation (Wash-out Tx). Stimulation of BCR signaling was done by the addition of 10 μg/mL anti-IgM for the last 15 min (+ ɑ-IgM). Whole-cell lysates were then analyzed for expression or phosphorylation of the indicated proteins (n=3 independent experiments per cell line). β-Tubulin serves as a loading control. Representative immunoblots are shown for each cell line.

Next, we sought to examine the effects of SRX3305 under activation induced BCR signaling. Following inhibitor treatment for 4 h, OSU-CLL and MEC-1 cells were stimulated with anti-IgM in the last 15 min. In parallel, inhibitor washout experiments were conducted to further evaluate whether inhibitor-induced target modulation could be maintained. CLL cells were subject to inhibitor treatment for 1 h prior to inhibitor washout. Three hours later, the cells were stimulated with anti-IgM (last 15 min) to elicit BCR crosslinking. In these studies, ibrutinib, a covalent irreversible BTK inhibitor, and the noncovalent reversible BTK inhibitor BMS-935177 were used as controls. As expected, continuous inhibitor treatment resulted in decreased phosphorylation of BTK and PRAS40 and reduced MYC expression in BCR-activated CLL cells. CLL cells treated with SRX3305 or ibrutinib (covalent BTK inhibitors) also retained inhibition of BTK phosphorylation following treatment washout, while cells treated with the reversible inhibitor BMS935177 regained BTK phosphorylation. Notably, SRX3305-treated cells maintained marked inhibition of PRAS40 phosphorylation and MYC expression after treatment washout (***Figure 2B***), whereas these effects were lost in ibrutinib-treated cells.

### 2.3 SRX3305 inhibits primary malignant B-cell survival and proliferation

To mimic conditions within pseudo-proliferation centers where CLL expands [40], we evaluated SRX3305 in primary CLL cells stimulated with CpG oligodeoxynucleotides (CpG ODN), a well-known toll-like receptor 9 agonist that promotes *ex vivo* CLL cell proliferation [41]. Characteristics of the patients’ samples used are summarized in ***Supplementary Table S2***. SRX3305 markedly decreased CpG ODN-induced proliferation of primary CLL cells in a dose-dependent manner (***Figure 3A***). Immunoblot analysis showed that SRX3305 reversed CpG ODN-mediated proliferation reflected by reduced MYC levels in primary CLL cells (***Figure 3B***). In parallel, SRX0035 induced the accumulation of P21 (cyclin-dependent kinase inhibitor), indicative of cell cycle arrest [42] (***Figure 3B***). SRX3305 did not influence proliferation of patient derived CLL cells under unstimulated/basal conditions (data not shown).

**Figure 3.**
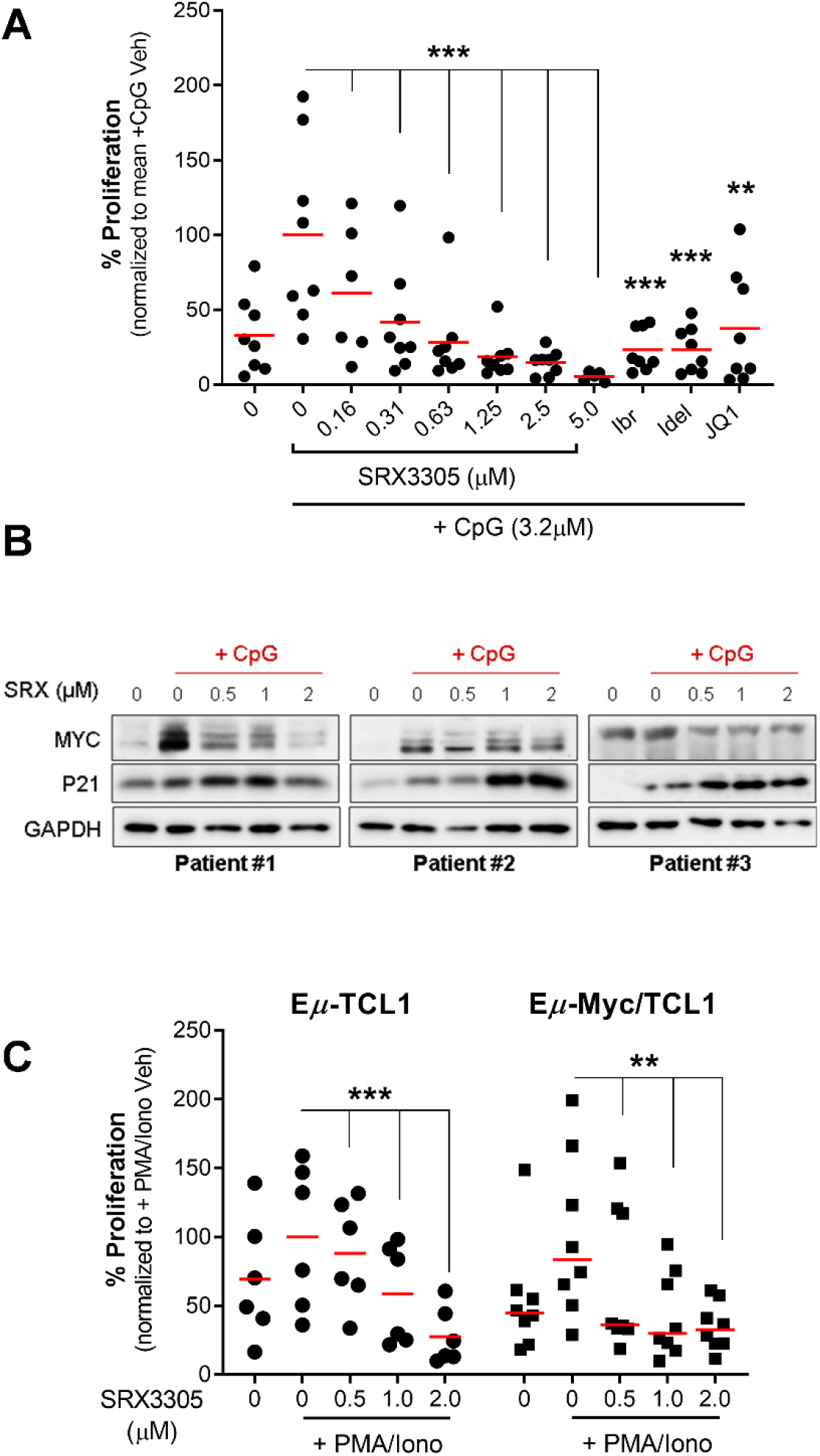
Antitumor effects of SRX3305 in primary CLL samples. **(A)** Patient-derived CLL cells (n=8) were treated with the indicated concentrations of inhibitors or DMSO vehicle and stimulated with CpG oligodeoxynucleotides (+ CpG; 3.2 μM) for 48 h. Proliferation was assessed via MTS assay and results are given as % proliferation normalized to mean stimulated vehicle (+ CpG Veh). Red lines indicate average values. *P* values indicate significance *vs*. stimulated vehicle. **(B)** Patient-derived CLL cells (n=6) were treated with DMSO vehicle or the indicated SRX3305 concentrations for 4 h in the presence of CpG oligonucleotides (+ CpG, 3.2 µM). The protein expression of MYC and P21 was determined by immunoblot analysis. GAPDH serves as a loading control. Representative immunoblots from three patients are shown. **(C)** Lymphocytes isolated from spleens of terminally diseased E*μ*-TCL1 (n=6, ●) or E*μ*-Myc/TCL1 (n=8, ■) mice were stimulated with 1X PMA/Ionomycin (+PMA/Iono) and treated with DMSO vehicle or increasing concentrations of SRX3305 as indicated. Proliferation was assessed via MTS assay 48 h later, and results are given as % proliferation normalized to mean stimulated vehicle (+PMA/Iono Veh). Red lines indicate averages. *P* values indicate significance *vs*. stimulated vehicle. ** *P*<0.01, *** *P*<0.001.

We next investigated SRX3305-induced cytotoxicity on tumor cells derived from murine models of aggressive B-cell leukemia and lymphoma. We utilized malignant B-cells isolated from E*μ*-TCL1 [43,44] (CLL model) and E*μ*-Myc/TCL1 mice [45,46] (concurrent CLL and B-cell lymphoma model). Primary murine spleen-derived tumor samples (comprising >90% malignant B-cells) were cultured *ex vivo* under mitogenic stimulation (PMA/ionomycin). SRX3305 treatment induced significant cytotoxicity in E*μ*-TCL1 and E*μ-*Myc/TCL1-derived malignant cells in a dose-dependent manner (***Figure 3C)***, suggesting promising therapeutic benefits in aggressive CLL and associated lymphomas (e.g., DLBCL-RT).

### 2.4 SRX3305 disrupts stroma survival support and chemokine-induced migration in CLL

Given the importance of the TME in CLL pathogenesis and therapy resistance [47-49], we sought to evaluate the efficacy of SRX3305 in the presence of stroma protection. The clinical efficacy of inhibitors targeting BCR signaling is partially attributed to disrupting CLL cell interactions with its TME [50]. As expected, co-culture of primary CLL cells on BM-derived stromal cells protects them from spontaneous apoptosis [49], resulting in a significant increase in CLL cell viability. Notably, treatment with SRX3305 reduced CLL cell viability in a dose-dependent manner despite stroma protection (***Figure 4A***). This was comparable to CLL therapeutics, ibrutinib or idelalisib. Of note, SRX3305 was not toxic to the stromal cells (***Supplementary Figure S2***), indicating that SRX3305-induced CLL cell cytotoxicity is not a function of reduced stroma viability.

**Figure 4.**
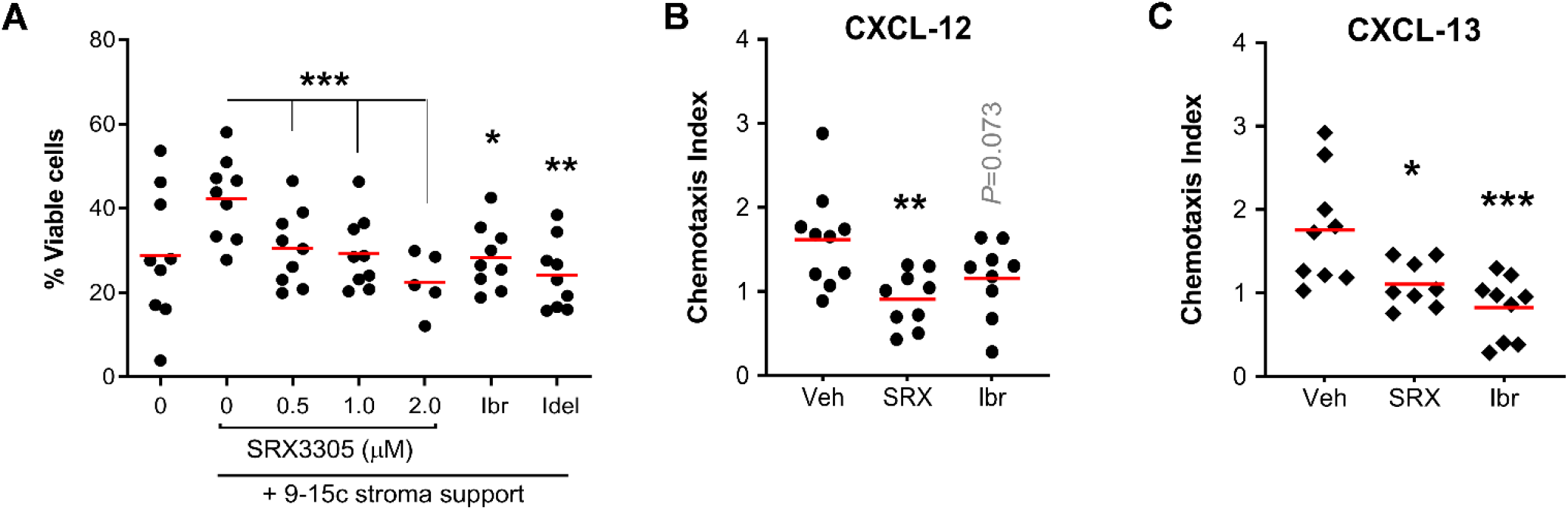
SRX3305 disrupts tumor microenvironment induced CLL cell survival and migration. Primary CLL cells (n=9 patients) were co-cultured with murine 9-15c bone marrow-derived stromal cells and incubated with increasing amounts of SRX3305 (0.5, 1, 2 µM), ibrutinib (Ibr, 1 µM), or idelalisib (Idel, 1 µM). After 48 h, CLL cell viability was determined by annexin V/PI staining, and cells negative for both annexin V and PI were declared to be viable. Red lines indicate average values. Significant dose-dependent decrease of CLL viability following SRX0035 treatment on stromal support is observed with asterisks denoting significance *vs*. vehicle control. **(B, C)** MEC-1 cells were treated with DMSO vehicle (Veh), SRX3305 (SRX, 1 µM) or ibrutinib (Ibr, 1 µM) and allowed to migrate for 6 h through trans-well inserts toward 200 ng/mL CXCl-12 (●, **B**) or 1000 ng/mL CXCL-13 (♦, **C**) Chemotaxis index represents the number of cells migrated toward the indicated chemokine divided by the number of cells migrated with no chemokine present for each treatment condition. Red lines indicate averages (n=10 independent experiments per chemokine). *P* values of inhibitor *vs*. vehicle are indicated. * *P*<0.05, ** *P*<0.01, *** *P*<0.001.

CLL cell migration, homing to supportive TME niches, and retention therein have been reported to support leukemic cell survival and disease progression [51,52]. We therefore investigated if SRX3305 could inhibit CLL cell migration towards key chemokines secreted by protective stromal cells in the BM (CXCL-12) and secondary lymphoid tissues (CXCL-13) [53-55]. Using a trans-well assay system, we demonstrate that SRX3305 reduced the migration of MEC-1 cells towards CXCL-12 or CXCL-13 (***Figure 4B – C***), further indicating that SRX3305 can disrupt CLL cell trafficking to supportive niches. Similar findings were seen in preclinical studies of the highly active BCR-targeting CLL therapeutic ibrutinib [4,56]. These findings suggest that SRX3305 can overcome TME-induced survival signaling, an established mediator in resistance to therapy in CLL.

### 2.5 SRX3305 is active in ibrutinib-resistant CLL

We hypothesized that because SRX3305 selectively targets PI3K and BRD4 in addition to BTK, this inhibitor would remain effective against ibrutinib-resistant CLL. To evaluate this in the context of non-genetic acquired drug resistance, we generated an ibrutinib-resistant CLL cell line model by prolonged culture of HG-3 cells with increasing amounts of ibrutinib (***Supplementary Figure S3***). Remarkably, SRX3305 decreased cell proliferation in ibrutinib-resistant HG3 (IR-HG3) cells (***Figure 5A***), while the anti-proliferative effects of ibrutinib were attenuated in IR-HG3 cells. These data suggest that SRX3305 has the potential to overcome acquired ibrutinib resistance. Further immunoblot analysis revealed that SRX3305 consistently decreased MYC expression and phosphorylation of PRAS (PI3K target), and increased P21 levels in IR-HG3 cells, while ibrutinib demonstrated a partial response and failed to reduce PRAS phosphorylation (***Figure 5B***). Overall BTK activation (phosphorylation) is reduced in the IR-HG3 cell line suggesting adaptive kinome reprogramming to bypass the effect of ibrutinib and an increased reliance on alternative survival mechanisms such as PI3K/AKT/ERK [7,57]. Hence, as expected we observed minimal decrease in BTK phosphorylation upon treatment with SRX3305 or ibrutinib.

**Figure 5.**
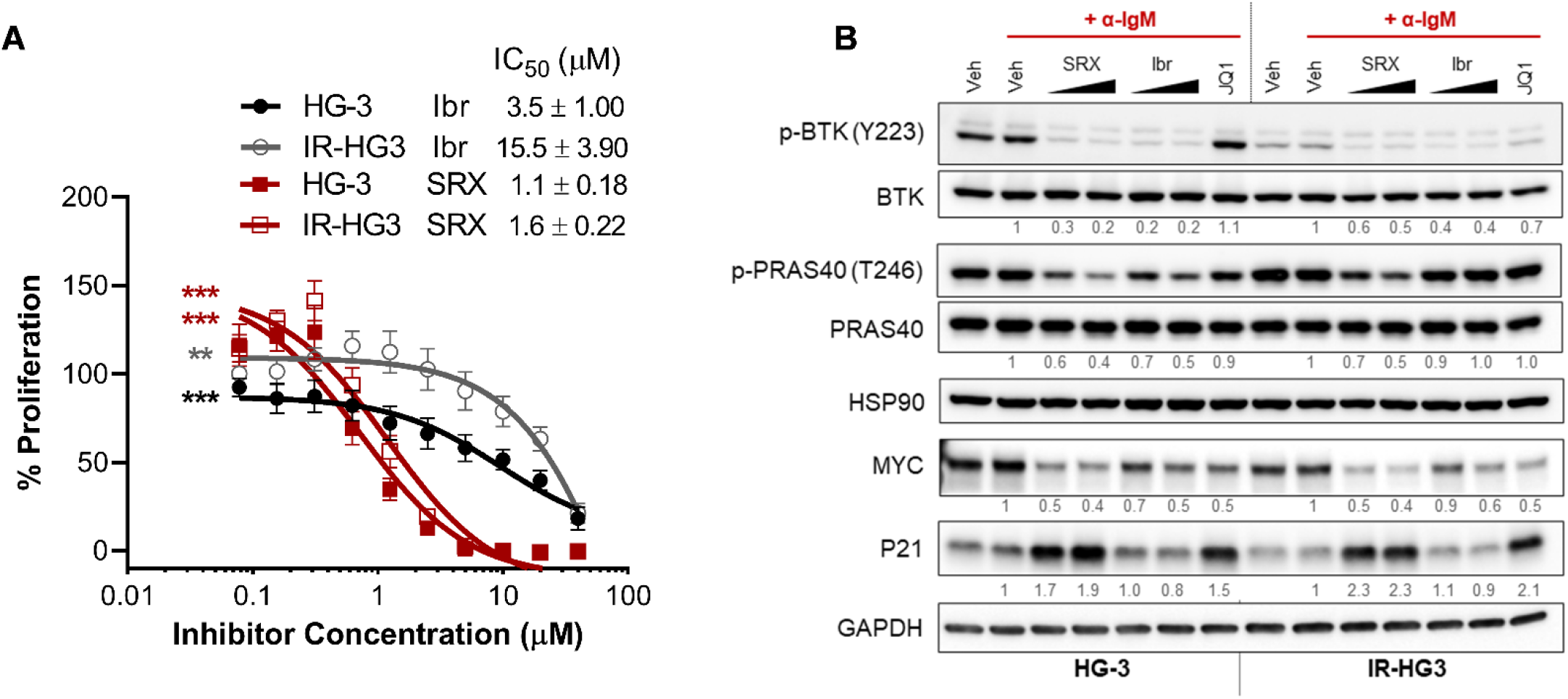
SRX3305 is effective in an ibrutinib-resistant CLL model. **(A)** HG-3 and ibrutinib-resistant HG-3 (IR-HG3) cells were incubated with increasing concentrations of ibrutinib or SRX3305 for 72 h. Proliferation was assessed via MTS assay and results are given as % proliferation normalized to vehicle. Error bars indicate SEM (n=6 independent experiments per cell line). Dose-dependent decrease in cell proliferation is observed with asterisks denoting significance. ** *P*<0.01, *** *P*<0.001. IC_50_ values are noted for each inhibitor within the in-figure legends (mean ± SEM). **(B)** Western blot analysis of HG-3 and ibrutinib-resistant HG-3 (IR-HG3) cells (n=3 independent experiments per cell line) that were treated with DMSO vehicle (Veh), SRX3305 (SRX; 1, 2 µM), or ibrutinib (Ibr; 1 µM) for 4 h and stimulated for the last 15 min with 10 µg/mL anti-IgM (+ ɑ-IgM). Representative immunoblots are shown for the phosphorylation and expression of the indicated proteins. Numbers below the bands represent densitometric quantification (relative to + ɑ-IgM vehicle). HSP90 or GAPDH were used as loading controls.

## 3. Discussion

Aberrant activities of diverse signaling pathways, including BTK, PI3K-AKT, and MYC-BRD4 contribute to CLL pathogenesis and persistence of residual disease with treatment. Each of these signaling pathways can be disrupted by the novel single small molecule BTK/PI3K/BRD4 inhibitor, SRX3305. In this study, we evaluated the preclinical efficacy of SRX3305 in CLL. We found that SRX3305 inhibits CLL cell proliferation at significantly lower doses than individual inhibitors of BTK, PI3K, and BRD4. SRX3305 proved to be not only cytostatic but also cytotoxic to CLL cells with marked induction of apoptosis. This response is in part due to irreversible binding of BTK, inhibiting its kinase function and thus significantly impairing BCR signaling. However, malignant B-cells can prioritize other signaling pathways to survive under consistent BTK inhibition. Importantly, SRX3305 also potently and irreversibly impairs phosphorylation of PI3K targets (e.g., PRAS40) and expression of BRD4 targets (e.g., MYC). Remarkably, SRX3305 sustained impressive anti-leukemic properties in ibrutinib-resistant CLL cells. SRX3305 also exhibited potent anti-tumor activity toward primary CLL cells *ex vivo*. The triple-inhibitor dose-dependently impaired the proliferation of stimulated primary CLL patient derived B-cells, and significantly reduced BTK phosphorylation and MYC expression in CLL cells while inducing P21 expression, indicative of cell cycle arrest. Lastly, SRX3305 significantly inhibited the proliferation of malignant B-cells from E*μ*-TCL1 and E*μ*-Myc/TCL1 mice *ex vivo*, suggesting promising therapeutic benefits in aggressive CLL and lymphomas.

CLL cells are highly dependent on diverse supportive stimuli produced by surrounding TME cells, including stromal cells in the BM and secondary lymphoid tissue niches [49]. Crosstalk between malignant and stromal cells in the TME can occur through direct cell-to-cell contact (via adhesion molecules) or indirectly by soluble factors (e.g., CXCL-12 and CXCL-13) [58]. These interactions lead to reciprocal activation of BCR and NF-κB pathways, gene expression changes (e.g., increase in MYC and anti-apoptotic proteins), and chemotaxis, resulting in sustained growth/proliferation of leukemic cells and resistance therapeutic agents [59]. In our study, primary CLL cells co-cultured with stromal cells maintained significantly greater viability *ex vivo*, and this was markedly reduced in the presence of SRX3305. Bidirectional signaling networks within the TME are integral to CLL disease progression and drug response. For instance, CXCL-12/CXCR4 and CXCL-13/CXCR5 are key chemokine networks that promote CLL cell homing to protective lymphoid tissues [53-55]. Inhibiting the BCR signaling pathway with ibrutinib [60] or idelalisib [61] is known to impair BCR- and chemokine-mediated cell adhesion and migration in CLL. Using a trans-well system, we show that treatment with SRX3305 significantly impairs CLL cell migration toward each CXCL-12 and CXCL-13. Preclinical studies of BET inhibitors in CLL [30,32] suggest that their efficacy may be partially attributed to interference with key CLL/TME interactions including those mediated by chemokine/cytokine networks. Interestingly, BRD4 was enriched at key CLL cell-trafficking genes (*CCR7, CXCR4*) in primary CLL cells [30]. Our preclinical findings demonstrate that SRX3305 can overcome TME-mediated survival signals suggesting its efficacy within TME sanctuaries in CLL.

Selective pressure of ibrutinib monotherapy often leads to the development of drug resistance in CLL. Ibrutinib can promote mutation of cysteine 481 of BTK, required for irreversible binding of ibrutinib to BTK. We previously demonstrated that SRX3262 and SRX3305 exhibited cytotoxic effects in malignant B-cells harboring the BTK-C481S mutation [37,38]. Besides mutations in drug-targeting proteins, several non-genetic mechanisms rendering CLL cells resistant to ibrutinib have been described and usually manifest as upregulation of alternative survival signaling pathways, such as the PI3K/AKT/ERK [57]. Here we evaluated SRX3305 in the context of acquired resistance to prolonged ibrutinib treatment. Notably, SRX3305 was almost equipotent in ibrutinib-resistant CLL cells and their parental ibrutinib-sensitive counterpart, whereas ibrutinib was significantly less cytotoxic in ibrutinib-resistant cells. Our ibrutinib-resistant CLL cells downregulate BTK activation, suggesting an increased reliance on alternative survival pathways. The phosphorylation of PI3K target PRAS40 was inhibited in ibrutinib-resistant and parental CLL cells following SRX3305 treatment. Similarly, MYC expression was equally downregulated in ibrutinib-resistant cells with SRX3305 treatment, illustrating the potential for a triple-inhibitor to bypass reciprocal activation of alternative survival signaling and overcome single-target drug resistance. These results suggest further evaluation of this novel chemotype in patients with CLL where responses to ibrutinib are partial or have relapse/refractory disease.

Numerous combination therapy approaches have been pursued for the treatment of CLL to combat emergence of drug resistance [14,15]. Preclinical studies have demonstrated that repressing MYC by targeting BET proteins enhances lymphoma cell vulnerability to PI3K inhibitors through upregulation of several PI3K pathways genes and increased GSK3β phosphorylation resulting in increased β-catenin protein abundance [62,63], suggesting that combinatory targeting of PI3K with SRX3305 would further promote MYC degradation by allowing GSK3β-dependent MYC phosphorylation [64] and stabilization of the MYC antagonist, MAD1 [65]. Our recent preclinical studies with the first-generation BTK/PI3K/BRD4 inhibitor, SRX3262 in MCL demonstrated reduced phosphorylation of MYC at Ser62 and Thr58 [37], critical mediators of MYC protein stability and degradation in cancer [64]. Furthermore, earlier studies have reported BET inhibition to be synergistically lethal with ibrutinib in MCL and CLL models [32,66]. PI3K and BTK inhibitors have also been combined to intercept constitutive BCR pathway signaling at multiple points [67]. While these approaches have shown promising preclinical efficacy, the increased risk of toxicity when combining multiple small molecule targeted agents remains an evident concern [38,68]. Herein lies the profound benefit of a single multi-target agent with the capability to simultaneously disrupt three key CLL pathogenic entities. A multi-target inhibitor could further allow simplification of dosing regimens for both patients and physicians. Future studies evaluating the pharmacokinetic and pharmacodynamic effects of this novel class of inhibitors *in vivo* are needed to delineate the translational potential of such promising TP-scaffold multi-action inhibitors for the treatment of B-cell malignancies.

## 4. Materials and Methods

### 4.1 Cell lines, primary samples, and inhibitors

Malignant B-cell lines (MEC-1, MEC-2, HG-3, OCI-LY3) were purchased from DSMZ (Braunschweig, Germany), while SU-DHL-6 cells were purchased from ATCC (Gaithersburg, MD). The OSU-CLL cell line [69] was provided by the Human Genetics Sample Bank of The Ohio State University (OSU; Columbus, OH). The murine 9-15c stromal cell line [70] was obtained from RIKEN (Ibaraki, Japan). To generate ibrutinib-resistant cells, HG-3 cells were treated with progressively increasing ibrutinib concentrations (up to 25 *μ*M) over the course of 3 months as further detailed in the ***Supplementary Materials***.

CLL patient samples (peripheral blood mononuclear cells; PBMCs) were obtained from the Leukemia Tissue Bank (OSU) in accordance with the Declaration of Helsinki following informed consent and under a protocol approved by the Institutional Review Board at OSU. In brief, PBMCs from CLL patients were isolated from whole blood through ficoll density gradient centrifugation. Prior to use in experiments, CLL patient-derived samples were confirmed to contain more than 90% CD5^+^/CD19^+^ cells by flow cytometric analysis on a NovoCyte 2060R flow cytometer (ACEA Biosciences Inc., San Diego, CA).

Lymphocytes isolated from E*μ*-TCL1 and E*μ*-Myc/TCL1 mouse spleens were obtained from the Animal Research Facility of the University of Nebraska Medical Center (UNMC), adhering to institutional animal care guidelines. Spleens were harvested from terminally ill mice (i.e., moribund at humane endpoints) and the percentage of malignant B-cells was confirmed via flow cytometry to exceed 90% CD5^+^/CD19^+^ prior to experimental use. Fluorochrome-conjugated CD19 and CD5 antibodies were ordered from BD Biosciences (Franklin Lakes, NJ).

Ibrutinib, idelalisib, JQ1 and OTX015 were purchased from Cayman Chemicals. SRX3262 and SRX3305 were synthesized and provided by SignalRx Pharmaceuticals, Inc. (Cumming, GA).

### 4.2 Cell culture

Cell lines and primary CLL samples were cultured in RPMI-1640 supplemented with 10% heat-inactivated fetal bovine serum (hi-FBS), 100 U/mL penicillin and 100 μg/mL streptomycin (P/S), and 2 μM L-glutamine except for OCI-LY3, which was maintained in Iscove’s modified Dulbecco’s medium (IMDM) supplemented with 20% hi-FBS, P/S, and 55 µM 2-mercaptoethanol. Primary murine splenocytes were cultured in RPMI-1640 supplemented with 10% hi-FBS, P/S, 2 mM L-glutamine, 55 µM 2-mercaptoethanol, 100 µM MEM non-essential amino acid solution, 1 mM sodium pyruvate, and 10 mM HEPES buffer. Hi-FBS was purchased from Avantor^®^ (Radnor, PA). Basal media and supplements were obtained from Life Technologies (Gaithersburg, MD).

### 4.3 Cytotoxicity and flow cytometric studies

MTS [3-(4,5-dimethylthiazol-2-yl)-5-(3-carboxymethoxyphenyl)-2-(4-sulfophenyl)-2H-tetrazolium] assays were used to determine inhibitor-induced cytotoxicity. Briefly, primary cells (∼0.7e^6^/well) or cell lines (∼25,000/well) were treated with vehicle (DMSO) or increasing inhibitor concentrations for up to 72 h in 96-well plates and then the CellTiter 96^®^ AQ_ueous_ assay (Promega, Madison, WI) was preformed according to manufacturer’s instruction to determine cell proliferation. Absorbance signal from each well was acquired at 490 nm on a Tecan Infinite^®^ M1000 Pro microplate reader (Männedorf, Switzerland). In addition, cell viability and/or apoptosis was measured by flow cytometry using annexin V/propidium iodide (PI) assay kit from Leinco Technologies (Fenton, MO) per manufacturer’s protocol. Stromal cell co-culture experiments were performed as previously described by plating a 75 cm^2^ flask (90% confluent) in 48-well plate 24 h before adding patient-derived CLL cells (1 e^7^/mL).

### 4.4 BCR pathway activation

To induce B-cell receptor crosslinking, OSU-CLL and MEC-1 were treated with 10 µg/mL of goat F(ab’)2 antihuman IgM (Jackson ImmunoResearch, West Grove, PA) for the final 15 min of treatment. Cells were then harvested and lysed for immunoblot analyses.

### 4.5 Inhibitor washout assay

OSU-CLL and MEC-1 cells were treated with indicated concentrations of inhibitors or vehicle for 1 h, washed three times with PBS, and incubated in complete culture medium for 3 h. For continuous treatment, the inhibitors or vehicle control remained for the entire treatment duration (4 h). In the final 15 min of treatment, cells were treated with anti-IgM to induce BCR crosslinking as described above and lysed for immunoblot analyses.

### 4.6 Ex vivo stimulation of primary malignant B-cells

Primary CLL cells were cultured with 3.2 *μ*M CpG 2006 oligodeoxynucleotides (Integrated DNA Technologies) to induce proliferation for the duration of treatment. For murine samples, 1X phorbol 12-myristate 13-acetate (PMA)/ionomycin cell stimulation cocktail (eBioscience, San Diego, CA) was added during treatments.

### 4.7 Immunoblot analyses

Cells were treated with indicated concentrations of inhibitors or vehicle control for 4 h under different stimuli. Thereafter, lysates were prepared, analyzed by sodium dodecyl sulfate polyacrylamide gel electrophoresis, and probed for select proteins. See ***Supplementary Methods*** for details.

### 4.8 Migration assay

Following 1 h pre-treatment with DMSO, SRX3305 or ibrutinib, MEC-1 cells were placed onto a 5-micron trans-well inserts (Corning, Tewksbury, MA) resting in wells containing 200 ng/mL CXCL-12 or 1000 ng/mL CXCL-13 (PeproTech, Cranbury, NJ). No chemokine control wells were included for each treatment condition. After incubating for 6 h, trans-well inserts were carefully removed, and the number of cells migrating through the insert towards the chemokines were counted by flow cytometry. Data was analyzed using NovoExpress software (ACEA Biosciences Inc., San Diego, CA).

### 4.9 Statistical analyses

Differences in cell viability and apoptosis between conditions of interest were assessed using ANOVA models. Dunnett’s test was used to compare conditions and dose levels with vehicle control. A test of linear trend for SRX3305 dosing was performed by employing orthogonal polynomial contrast coefficients. Log-transformations of the data were applied, when necessary, before modeling to stabilize variances specifically for data analyzed as raw absorption. Percent proliferation data or percent viable data was analyzed without transformation. Data is represented as mean ± SEM. Analyses were done using SAS/STAT software, version 9.4 of the SAS System for Windows (SAS Institute Inc., Cary, NC). P values < 0.05 were considered statistically significant.

## 5. Conclusion

The novel BTK/PI3K/BRD4 inhibitor, SRX3305 demonstrates marked anti-tumor properties in preclinical models of CLL. Importantly, the pro-survival, proliferative, therapy-resistant, and activation effects promoted by TME sanctuaries were abrogated by SRX3305, which furthermore blocked CLL cell chemotaxis. Additionally, SRX3305 sustained impressive anti-tumor effects in ibrutinib-resistant CLL cells and effectively attenuated alternative/downstream survival signaling pathways. Together, these findings provide strong rationale for the clinical development of this distinct triple-action inhibitor for treating B-cell lymphoproliferative diseases, especially in the relapse/refractory setting.

## Acknowledgments

We thank SignalRx Pharmaceuticals, Inc for providing SRX3305 and SRX3262 and Dr. Donald L. Durden for discussion. The authors also thank The Ohio State University Leukemia Tissue Bank staff for primary patient sample procurement services that was supported by the National Cancer Institute Grant P30 CA016058.

## Supplementary Materials

The following supporting information can be downloaded at:

## Author Contributions

A.L.S., A.P.E., D.Y.M., L.T.U. and D.E. performed experiments, and acquired and/or analyzed the data. L.M.S. and E.R.L assisted with the statistical analysis. A.L.S., T.G.K, and D.E. contributed to the conceptualization, experimental design, and interpretation of the results. A.L.S. and D.E. wrote the original manuscript draft and obtained input from all other authors who reviewed and edited the manuscript. D.E. managed and supervised the study and obtained funding. All authors have read and agreed to the published version of the manuscript.

## Funding

This work was supported by institutional funds from the University of Nebraska Medical Center (D.E.).

## Institutional Review Board Statement

Primary PBMCs from CLL patients were obtained from the Leukemia Tissue Bank at The Ohio State University Comprehensive Cancer Center in accordance with the Declaration of Helsinki following informed consent under approved Institutional Review Board protocol (#1997C0194). The samples were de-identified and used for the *in vitro* experiments outlined in this study. Primary murine tumor samples were obtained through an approved institutional animal care and use committee protocol at UNMC (#1816901FC).

## Informed Consent Statement

Patients provided informed consent for the use of their biologic specimens for research purposes according to institutional guidelines.

## Data Availability Statement

The structures of SRX3305 or SRX3260 used in this study are proprietary information of SignalRx Pharmaceuticals, Inc. (*Patent number WO2020023340A1*, 2020) [36] and are not publicly available. All other data will be made available from the authors upon reasonable request.

## Conflicts of Interest

A.L.S., A.P.E., D.Y.M., L.T.U., L.M.S., E.R.L., A.K., T.G.K., and D.E. have no conflicts of interest. C.R.D. has served on an advisory board for TG Therapeutics. J.M.V. has received honoraria from AbbVie, Janssen Pharmaceuticals, AstraZeneca, MEI Pharma, Johnson & Johnson, Pharmacyclics, Genentech, Inc., MorphoSys, and Lilly.

## Notes

### Summary of Updates

Figure 2 labeling revised.

## References

1. Kipps TJ, Stevenson FK, Wu CJ, Croce CM, Packham G, Wierda WG, et al. Chronic lymphocytic leukaemia. Nat Rev Dis Primers. 2017;3:17008.10.1038/nrdp.2017.8

2. Riches JC, Gribben JG. Understanding the immunodeficiency in chronic lymphocytic leukemia: potential clinical implications. Hematol Oncol Clin North Am. 2013;27(2):207–235.10.1016/j.hoc.2013.01.003

3. Hallek M. Chronic lymphocytic leukemia: 2020 update on diagnosis, risk stratification and treatment. Am J Hematol. 2019;94(11):1266–1287.10.1002/ajh.25595

4. Herman SE, Gordon AL, Hertlein E, Ramanunni A, Zhang X, Jaglowski S, et al. Bruton tyrosine kinase represents a promising therapeutic target for treatment of chronic lymphocytic leukemia and is effectively targeted by PCI-32765. Blood. 2011;117(23):6287–6296.10.1182/blood-2011-01-328484

5. Herman SM JJ, Mustafa RZ, Farooqui M, Wiestner A. In Vivo Effects Of Ibrutinib On The Migration Of Chronic Lymphocytic Leukemia Cells Differ Between Patients and Reduce The Ability Of The Bone Marrow Microenvironment To Attract The Tumor Cells. Blood. 2013;122(21):604

6. O’Brien S, Furman RR, Coutre S, Flinn IW, Burger JA, Blum K, et al. Single-agent ibrutinib in treatmentnaive and relapsed/refractory chronic lymphocytic leukemia: a 5-year experience. Blood. 2018;131(17):1910–1919.10.1182/blood-2017-10-810044

7. Zhao X, Lwin T, Silva A, Shah B, Tao J, Fang B, et al. Unification of de novo and acquired ibrutinib resistance in mantle cell lymphoma. Nat Commun. 2017;8:14920.10.1038/ncomms14920

8. Rahal R, Frick M, Romero R, Korn JM, Kridel R, Chan FC, et al. Pharmacological and genomic profiling identifies NF-kappaB-targeted treatment strategies for mantle cell lymphoma. Nat Med. 2014;20(1):87–92.10.1038/nm.3435

9. Moyo TK, Wilson CS, Moore DJ, Eischen CM. Myc enhances B-cell receptor signaling in precancerous B cells and confers resistance to Btk inhibition. Oncogene. 2017;36(32):4653–4661.10.1038/onc.2017.95

10. Lee J, Zhang LL, Wu W, Guo H, Li Y, Sukhanova M, et al. Activation of MYC, a bona fide client of HSP90, contributes to intrinsic ibrutinib resistance in mantle cell lymphoma. Blood Adv. 2018;2(16):2039–2051.10.1182/bloodadvances.2018016048

11. Ahn IE, Underbayev C, Albitar A, Herman SE, Tian X, Maric I, et al. Clonal evolution leading to ibrutinib resistance in chronic lymphocytic leukemia. Blood. 2017;129(11):1469–1479.10.1182/blood-2016-06-719294

12. George B, Chowdhury SM, Hart A, Sircar A, Singh SK, Nath UK, et al. Ibrutinib Resistance Mechanisms and Treatment Strategies for B-Cell lymphomas. Cancers (Basel). 2020;12(5).10.3390/cancers12051328

13. Rossi D, Spina V, Gaidano G. Biology and treatment of Richter syndrome. Blood. 2018;131(25):2761–2772.10.1182/blood-2018-01-791376

14. Skanland SS, Mato AR. Overcoming resistance to targeted therapies in chronic lymphocytic leukemia. Blood Adv. 2021;5(1):334–343.10.1182/bloodadvances.2020003423

15. Timofeeva N, Gandhi V. Ibrutinib combinations in CLL therapy: scientific rationale and clinical results. Blood Cancer J. 2021;11(4):79.10.1038/s41408-021-00467-7

16. Baylin SB, Jones PA. A decade of exploring the cancer epigenome - biological and translational implications. Nat Rev Cancer. 2011;11(10):726–734.10.1038/nrc3130

17. Mansouri L, Wierzbinska JA, Plass C, Rosenquist R. Epigenetic deregulation in chronic lymphocytic leukemia: Clinical and biological impact. Semin Cancer Biol. 2018;51:1–11.10.1016/j.semcancer.2018.02.001

18. Xanthopoulos C, Kostareli E. Advances in Epigenetics and Epigenomics in Chronic Lymphocytic Leukemia. Current Genetic Medicine Reports. 2019;7(4):214–226.10.1007/s40142-019-00178-3

19. Jiang Y, Redmond D, Nie K, Eng KW, Clozel T, Martin P, et al. Deep sequencing reveals clonal evolution patterns and mutation events associated with relapse in B-cell lymphomas. Genome Biol. 2014;15(8):432.10.1186/s13059-014-0432-0

20. Filippakopoulos P, Knapp S. Targeting bromodomains: epigenetic readers of lysine acetylation. Nature Reviews Drug Discovery. 2014;13(5):337–356.10.1038/nrd4286

21. Hnisz D, Abraham BJ, Lee TI, Lau A, Saint-Andre V, Sigova AA, et al. Super-enhancers in the control of cell identity and disease. Cell. 2013;155(4):934–947.10.1016/j.cell.2013.09.053

22. Loven J, Hoke HA, Lin CY, Lau A, Orlando DA, Vakoc CR, et al. Selective inhibition of tumor oncogenes by disruption of super-enhancers. Cell. 2013;153(2):320–334.10.1016/j.cell.2013.03.036

23. Chapuy B, McKeown MR, Lin CY, Monti S, Roemer MG, Qi J, et al. Discovery and characterization of super-enhancer-associated dependencies in diffuse large B cell lymphoma. Cancer Cell. 2013;24(6):777–790.10.1016/j.ccr.2013.11.003

24. Dawson MA, Prinjha RK, Dittmann A, Giotopoulos G, Bantscheff M, Chan WI, et al. Inhibition of BET recruitment to chromatin as an effective treatment for MLL-fusion leukaemia. Nature. 2011;478(7370):529–533.10.1038/nature10509

25. Hogg SJ, Newbold A, Vervoort SJ, Cluse LA, Martin BP, Gregory GP, et al. BET Inhibition Induces Apoptosis in Aggressive B-Cell Lymphoma via Epigenetic Regulation of BCL-2 Family Members. Mol Cancer Ther. 2016;15(9):2030–2041.10.1158/1535-7163.MCT-15-0924

26. Huang B, Yang XD, Zhou MM, Ozato K, Chen LF. Brd4 coactivates transcriptional activation of NF-kappaB via specific binding to acetylated RelA. Mol Cell Biol. 2009;29(5):1375–1387.10.1128/MCB.01365-08

27. Mansouri L, Papakonstantinou N, Ntoufa S, Stamatopoulos K, Rosenquist R. NF-kappaB activation in chronic lymphocytic leukemia: A point of convergence of external triggers and intrinsic lesions. Semin Cancer Biol. 2016;39:40–48.10.1016/j.semcancer.2016.07.005

28. Chaidos A, Caputo V, Karadimitris A. Inhibition of bromodomain and extra-terminal proteins (BET) as a potential therapeutic approach in haematological malignancies: emerging preclinical and clinical evidence. Ther Adv Hematol. 2015;6(3):128–141.10.1177/2040620715576662

29. Stathis A, Bertoni F. BET Proteins as Targets for Anticancer Treatment. Cancer Discov. 2018;8(1):24–36.10.1158/2159-8290.CD-17-0605

30. Ozer HG, El-Gamal D, Powell B, Hing ZA, Blachly JS, Harrington B, et al. BRD4 Profiling Identifies Critical Chronic Lymphocytic Leukemia Oncogenic Circuits and Reveals Sensitivity to PLX51107, a Novel Structurally Distinct BET Inhibitor. Cancer Discov. 2018;8(4):458–477.10.1158/2159-8290.CD-17-0902

31. Spriano F, Stathis A, Bertoni F. Targeting BET bromodomain proteins in cancer: The example of lymphomas. Pharmacol Ther. 2020;215:107631.10.1016/j.pharmthera.2020.107631

32. Kim E, Ten Hacken E, Sivina M, Clarke A, Thompson PA, Jain N, et al. The BET inhibitor GS-5829 targets chronic lymphocytic leukemia cells and their supportive microenvironment. Leukemia. 2020;34(6):1588–1598.10.1038/s41375-019-0682-7

33. Carra G, Nicoli P, Lingua MF, Maffeo B, Cartella A, Circosta P, et al. Inhibition of bromodomain and extra-terminal proteins increases sensitivity to venetoclax in chronic lymphocytic leukaemia. J Cell Mol Med. 2020;24(2):1650–1657.10.1111/jcmm.14857

34. Tarantelli C, Lange M, Gaudio E, Cascione L, Spriano F, Kwee I, et al. Copanlisib synergizes with conventional and targeted agents including venetoclax in B-and T-cell lymphoma models. Blood Adv. 2020;4(5):819–829.10.1182/bloodadvances.2019000844

35. Fiskus W, Mill CP, Perera D, Birdwell C, Deng Q, Yang H, et al. BET proteolysis targeted chimera-based therapy of novel models of Richter Transformation-diffuse large B-cell lymphoma. Leukemia. 2021;35(9):2621–2634.10.1038/s41375-021-01181-w

36. Morales GA, Garlich JR, Durden DL, Inventors. Single molecule compounds providing multi-target inhibition of btk and other proteins and methods of use thereof. 2020.

37. Pal D, Vann KR, Joshi S, Sahar NE, Morales GA, El-Gamal D, et al. The BTK/PI3K/BRD4 axis inhibitor SRX3262 overcomes Ibrutinib resistance in mantle cell lymphoma. iScience. 2021;24(9):102931.10.1016/j.isci.2021.102931

38. Vann KR, Pal D, Smith AL, Sahar NE, Krishnaiah M, El-Gamal D, et al. Combinatorial inhibition of BTK, PI3K-AKT and BRD4-MYC as a strategy for treatment of mantle cell lymphoma. Mol Biomed. 2022;3(1):2.10.1186/s43556-021-00066-9

39. Seda V, Mraz M. B-cell receptor signalling and its crosstalk with other pathways in normal and malignant cells. Eur J Haematol. 2015;94(3):193–205.10.1111/ejh.12427

40. Swerdlow SH, Murray LJ, Habeshaw JA, Stansfeld AG. Lymphocytic lymphoma/B-chronic lymphocytic leukaemia--an immunohistopathological study of peripheral B lymphocyte neoplasia. Br J Cancer. 1984;50(5):587–599.10.1038/bjc.1984.225

41. Decker T, Schneller F, Sparwasser T, Tretter T, Lipford GB, Wagner H, et al. Immunostimulatory CpG-oligonucleotides cause proliferation, cytokine production, and an immunogenic phenotype in chronic lymphocytic leukemia B cells. Blood. 2000;95(3):999–1006

42. Abbas T, Dutta A. p21 in cancer: intricate networks and multiple activities. Nat Rev Cancer. 2009;9(6):400–414.10.1038/nrc2657

43. Bichi R, Shinton SA, Martin ES, Koval A, Calin GA, Cesari R, et al. Human chronic lymphocytic leukemia modeled in mouse by targeted TCL1 expression. Proc Natl Acad Sci U S A. 2002;99(10):6955–6960.10.1073/pnas.102181599

44. Johnson AJ, Lucas DM, Muthusamy N, Smith LL, Edwards RB, De Lay MD, et al. Characterization of the TCL-1 transgenic mouse as a preclinical drug development tool for human chronic lymphocytic leukemia. Blood. 2006;108(4):1334–1338.10.1182/blood-2005-12-011213

45. Rogers KA, El-Gamal D, Bonnie HK, Zachary HA, Virginia GM, Rose M, et al. The Eµ-Myc/TCL1 Transgenic Mouse As a New Aggressive B-Cell Malignancy Model Suitable for Preclinical Therapeutics Testing. Blood. 2015;126(23):2752–2752.10.1182/blood.V126.23.2752.2752

46. Lucas F, Rogers KA, Harrington BK, Pan A, Yu L, Breitbach J, et al. Emu-TCL1xMyc: A Novel Mouse Model for Concurrent CLL and B-Cell Lymphoma. Clin Cancer Res. 2019;25(20):6260–6273.10.1158/1078-0432.CCR-19-0273

47. Herishanu Y, Katz BZ, Lipsky A, Wiestner A. Biology of chronic lymphocytic leukemia in different microenvironments: clinical and therapeutic implications. Hematol Oncol Clin North Am. 2013;27(2):173–206.10.1016/j.hoc.2013.01.002

48. Ramsay AD, Rodriguez-Justo M. Chronic lymphocytic leukaemia--the role of the microenvironment pathogenesis and therapy. Br J Haematol. 2013;162(1):15–24.10.1111/bjh.12344

49. Kurtova AV, Balakrishnan K, Chen R, Ding W, Schnabl S, Quiroga MP, et al. Diverse marrow stromal cells protect CLL cells from spontaneous and drug-induced apoptosis: development of a reliable and reproducible system to assess stromal cell adhesion-mediated drug resistance. Blood. 2009;114(20):4441–4450.10.1182/blood-2009-07-233718

50. Burger JA, Wiestner A. Targeting B cell receptor signalling in cancer: preclinical and clinical advances. Nat Rev Cancer. 2018;18(3):148–167.10.1038/nrc.2017.121

51. Hartmann TN, Grabovsky V, Wang W, Desch P, Rubenzer G, Wollner S, et al. Circulating B-cell chronic lymphocytic leukemia cells display impaired migration to lymph nodes and bone marrow. Cancer Res. 2009;69(7):3121–3130.10.1158/0008-5472.CAN-08-4136

52. Scielzo C, Bertilaccio MT, Simonetti G, Dagklis A, ten Hacken E, Fazi C, et al. HS1 has a central role in the trafficking and homing of leukemic B cells. Blood. 2010;116(18):3537–3546.10.1182/blood-2009-12-258814

53. Burger JA, Burger M, Kipps TJ. Chronic lymphocytic leukemia B cells express functional CXCR4 chemokine receptors that mediate spontaneous migration beneath bone marrow stromal cells. Blood. 1999;94(11):3658–3667

54. Ma Q, Jones D, Springer TA. The chemokine receptor CXCR4 is required for the retention of B lineage and granulocytic precursors within the bone marrow microenvironment. Immunity. 1999;10(4):463–471.10.1016/s1074-7613(00)80046-1

55. Ansel KM, Harris RB, Cyster JG. CXCL13 is required for B1 cell homing, natural antibody production, and body cavity immunity. Immunity. 2002;16(1):67–76.10.1016/s1074-7613(01)00257-6

56. Ponader S, Chen SS, Buggy JJ, Balakrishnan K, Gandhi V, Wierda WG, et al. The Bruton tyrosine kinase inhibitor PCI-32765 thwarts chronic lymphocytic leukemia cell survival and tissue homing in vitro and in vivo. Blood. 2012;119(5):1182–1189.10.1182/blood-2011-10-386417

57. Forestieri G, Terzi di Bergamo L, Loh J, Spina V, Zucchetto A, Condoluci A. Mechanisms Of Adaptation To Ibrutinib In High Risk Chronic Lymphocytic Leukemia. Paper presented at:25th Congress of the European Hematology Association 2020; 294974; S154.

58. Dubois N, Crompot E, Meuleman N, Bron D, Lagneaux L, Stamatopoulos B. Importance of Crosstalk Between Chronic Lymphocytic Leukemia Cells and the Stromal Microenvironment: Direct Contact, Soluble Factors, and Extracellular Vesicles. Front Oncol. 2020;10:1422.10.3389/fonc.2020.01422

59. Shain KH, Dalton WS, Tao J. The tumor microenvironment shapes hallmarks of mature B-cell malignancies. Oncogene. 2015;34(36):4673–4682.10.1038/onc.2014.403

60. de Rooij MF, Kuil A, Geest CR, Eldering E, Chang BY, Buggy JJ, et al. The clinically active BTK inhibitor PCI-32765 targets B-cell receptor-and chemokine-controlled adhesion and migration in chronic lymphocytic leukemia. Blood. 2012;119(11):2590–2594.10.1182/blood-2011-11-390989

61. Hoellenriegel J, Meadows SA, Sivina M, Wierda WG, Kantarjian H, Keating MJ, et al. The phosphoinositide 3’-kinase delta inhibitor, CAL-101, inhibits B-cell receptor signaling and chemokine networks in chronic lymphocytic leukemia. Blood. 2011;118(13):3603–3612.10.1182/blood-2011-05-352492

62. Chen ZQ, Cao ZR, Wang Y, Zhang X, Xu L, Wang YX, et al. Repressing MYC by targeting BET synergizes with selective inhibition of PI3Kalpha against B cell lymphoma. Cancer Lett. 2022;524:206–218.10.1016/j.canlet.2021.10.022

63. Derenzini E, Mondello P, Erazo T, Portelinha A, Liu Y, Scallion M, et al. BET Inhibition-Induced GSK3beta Feedback Enhances Lymphoma Vulnerability to PI3K Inhibitors. Cell Rep. 2018;24(8):2155–2166.10.1016/j.celrep.2018.07.055

64. Sears R, Nuckolls F, Haura E, Taya Y, Tamai K, Nevins JR. Multiple Ras-dependent phosphorylation pathways regulate Myc protein stability. Genes Dev. 2000;14(19):2501–2514.10.1101/gad.836800

65. Zhu J, Blenis J, Yuan J. Activation of PI3K/Akt and MAPK pathways regulates Myc-mediated transcription by phosphorylating and promoting the degradation of Mad1. Proc Natl Acad Sci U S A. 2008;105(18):6584–6589.10.1073/pnas.0802785105

66. Schaffer M, Chaturvedi S, Davis C, Aquino R, Stepanchick E, Versele M, et al. Identification of potential ibrutinib combinations in hematological malignancies using a combination high-throughput screen. Leuk Lymphoma. 2018;59(4):931–940.10.1080/10428194.2017.1349899

67. Niemann CU, Mora-Jensen HI, Dadashian EL, Krantz F, Covey T, Chen SS, et al. Combined BTK and PI3Kdelta Inhibition with Acalabrutinib and ACP-319 Improves Survival and Tumor Control in CLL Mouse Model. Clin Cancer Res. 2017;23(19):5814–5823.10.1158/1078-0432.CCR-17-0650

68. Andrews FH, Singh AR, Joshi S, Smith CA, Morales GA, Garlich JR, et al. Dual-activity PI3K-BRD4 inhibitor for the orthogonal inhibition of MYC to block tumor growth and metastasis. Proc Natl Acad Sci U S A. 2017;114(7):E1072–E1080.10.1073/pnas.1613091114

69. Hertlein E, Beckwith KA, Lozanski G, Chen TL, Towns WH, Johnson AJ, et al. Characterization of a new chronic lymphocytic leukemia cell line for mechanistic in vitro and in vivo studies relevant to disease. PLoS One. 2013;8(10):e76607.10.1371/journal.pone.0076607

70. Yamada Y, Sakurada K, Takeda Y, Gojo S, Umezawa A. Single-cell-derived mesenchymal stem cells overexpressing Csx/Nkx2.5 and GATA4 undergo the stochastic cardiomyogenic fate and behave like transient amplifying cells. Exp Cell Res. 2007;313(4):698–706.10.1016/j.yexcr.2006.11.012

71. Schneider CA, Rasband WS, Eliceiri KW. NIH Image to ImageJ: 25 years of image analysis. Nat Methods. 2012;9(7):671–675.10.1038/nmeth.2089

72. Rasul E, Salamon D, Nagy N, Leveau B, Banati F, Szenthe K, et al. The MEC1 and MEC2 lines represent two CLL subclones in different stages of progression towards prolymphocytic leukemia. PLoS One. 2014;9(8):e106008.10.1371/journal.pone.0106008

73. Stacchini A, Aragno M, Vallario A, Alfarano A, Circosta P, Gottardi D, et al. MEC1 and MEC2: two new cell lines derived from B-chronic lymphocytic leukaemia in prolymphocytoid transformation. Leuk Res. 1999;23(2):127–136.10.1016/s0145-2126(98)00154-4

74. Rosen A, Bergh AC, Gogok P, Evaldsson C, Myhrinder AL, Hellqvist E, et al. Lymphoblastoid cell line with B1 cell characteristics established from a chronic lymphocytic leukemia clone by in vitro EBV infection. Oncoimmunology. 2012;1(1):18–27.10.4161/onci.1.1.18400

75. Mehra S, Messner H, Minden M, Chaganti RS. Molecular cytogenetic characterization of non-Hodgkin lymphoma cell lines. Genes Chromosomes Cancer. 2002;33(3):225–234.10.1002/gcc.10025

76. Quentmeier H, Pommerenke C, Dirks WG, Eberth S, Koeppel M, MacLeod RAF, et al. The LL-100 panel: 100 cell lines for blood cancer studies. Sci Rep. 2019;9(1):8218.10.1038/s41598-019-44491-x

77. Drexler HG, Quentmeier H. The LL-100 Cell Lines Panel: Tool for Molecular Leukemia-Lymphoma Research. Int J Mol Sci. 2020;21(16).10.3390/ijms21165800

78. Davis RE, Ngo VN, Lenz G, Tolar P, Young RM, Romesser PB, et al. Chronic active B-cell-receptor signalling in diffuse large B-cell lymphoma. Nature. 2010;463(7277):88–92.10.1038/nature08638

79. Ngo VN, Young RM, Schmitz R, Jhavar S, Xiao W, Lim KH, et al. Oncogenically active MYD88 mutations in human lymphoma. Nature. 2011;470(7332):115–119.10.1038/nature09671

80. Morin RD, Johnson NA, Severson TM, Mungall AJ, An J, Goya R, et al. Somatic mutations altering EZH2 (Tyr641) in follicular and diffuse large B-cell lymphomas of germinal-center origin. Nat Genet. 2010;42(2):181–185.10.1038/ng.518

