## Supplemental Methods and Figures for "A novel triple-action inhibitor targeting B-cell receptor signaling and BRD4 demonstrates preclinical activity in chronic lymphocytic leukemia"

### Supplementary Materials

### Supplementary Methods

#### Immunoblot analysis

Samples were lysed in RIPA buffer containing Sigma Protease and Phosphatase Inhibitor Cocktails and phenylmethyl sulfonyl fluoride (Sigma-Aldrich, Saint Louis, MO), and analyzed by sodium dodecyl sulfate polyacrylamide gel electrophoresis (SDS-PAGE). Gels were transferred onto nitrocellulose membranes using Bio-Rad's TransBlot Turbo Transfer System, probed with primary antibodies and HRP-conjugated secondary antibodies (Bio-Rad Laboratories, Hercules, CA). Blots were then visualized on Bio-Rad's ChemiDoc MP imager using SuperSignal West Pico PLUS Chemiluminescent Substrate or Femto Maximum Sensitivity Substrate (Thermo Fisher Scientific, Waltham, MA) according to the manufacturer's instructions. ImageJ software was used for densitometric band quantification [71].

Primary antibodies against phospho-BTK (Tyr223), BTK, phospho-PRAS40 (Thr246), PRAS40, phospho-ERK (Thr202/Tyr204), ERK, MYC, P21, and GAPDH were obtained from Cell Signaling Technologies (Danvers, MA). Anti-β-TUBULIN was ordered from Sigma-Aldrich and anti-HSP 90α/β was obtained from Santa Cruz Biotechnologies (Santa Cruz, CA).

### Supplementary Data

### Supplementary Tables

### Table S1. Key characteristics of the malignant B-cell lines used in the study

| Cell Line | Disease Type | Key Characteristics |
|---|---|---|
| MEC-1 [1] | CLL | Mutated IGHV, 17p deletion, complex karyotype [2] |
| MEC-2 [1] | CLL | Mutated IGHV, 17p deletion, complex karyotype |
| OSU-CLL[3] | CLL | Mutated IGHV, non-complex karyotype |
| HG-3 [4] | CLL | Un-mutated IGHV, non-complex karyotype |
| OCI-LY3 [5] | ABC-DLBCL | t(14;19)(q32;q13) IGH-SPIB, amplified BCL2<br>MYD88 mut, CARD11 mut |
| SU-DHL-6 [6] | GC-DLBCL | t(14;18)(q32;q21), IGH-BCL2, rearranged BCL6 gene, EZH2 mut |

1 MEC-1 and MEC-2 cell lines represent spontaneous outgrowths of CLL clones on two subsequent occasions (1 y apart) from a patient with mutated IGHV CLL and a complex karyotype with 17p deletion ($P53^{mutated}$). MEC-2 cells have evident prolymphocytic leukemia transformation [72,73].

2 Complex karyotype is defined as having ≥3 chromosomal aberrations [72].

3 The OSU-CLL cell line originated from a CLL patient with mutated IGHV and non-complex karyotype [69].

4 The HG-3 cell line originated from a CLL patient with un-mutated IGHV and demonstrates non-complex karyotype with biallelic 13q14 deletions [74].

5 The activated B-cell like DLBCL (ABC-DLBCL) cell line, OCI-LY3 carries cryptic t(14;19) with rearrangement of IGH and SPIB and contains copy number amplification of the BCL2 region [75-77]. OCI-LY3 cells harbor mutations (mut) in CARD11 and MYD88 (L265P) [78,79].

6 The germinal center-like DLBCL (GC-DLBCL) cell line, SU-DHL-6 carries t(14:18) effecting IGH-BCL2 fusion and has BCL6 gene rearrangement [76,77]. SU-DHL-6 cells harbor the EZH2 Y641N mutation [80].

### Table S2. Characteristics of CLL patient samples used in the study

| ID | Gender | Age | IGHV status | Treatment status | Karyotype | Figure(s) |
|---|---|---|---|---|---|---|
| 6-0615 | F | 65 | U | N | 47,XX,+12(1)/46,XX(19) | 3A, 3B (patient #1*) |
| 6-0726 | M | 58 | U | N | 46,XY(23)/nonclonal(4) | 3A, 3B (patient #2*) |
| 12-1833 | M | 76 | U | T | 44,XY,t(1;19)(q21;p13.1),-4,der(8;18)(q10;10),-17,+der(?)t(?;4)(?;q12)[cp10]/44,sl,add(1)(p36.1),add(9)(q31)[cp4]/44,sl,t(1;22)(p32;p11.2),-der(8;18),+der(8;18)(q10;q10)t(?;8)(?;q24)t(?;4)(?;q12),+mar[cp3]/43-45,XY,-4,t(5;16)(q13;p13.3),del(7)(q32),-17,add(22)(p11.2),+der(?)t(?;4)(?;q12)[cp4] | 3B (patient #3*), 4A |
| 6-1127 | F | 76 | U | N | 46,XX,del(11)(q13q23)(6)/45,X,-X(3)/46,XX(10)/nonclonal | 3A, 3B |
| 16-2100 | F | 79 | U | T | 46,XY,der(1)(11qter->q13::?::1p13->1qter),del(7)(q32q36),-11,der(20)t(15;20)(q15;q13.3),+mar[3,one w/ nonclonal abnormalities]/ 46,sl,add(8)(p11.2)[16,one w/ nonclonal abnormalities]/non-clonal[1] | 3B |
| 18-4770 | M | 50 | U | N | 47,XY,+12,t(14;19)(q32.3;q13.2)[10]/46,XY[10] | 3A, 3B |
| 8-1148 | F | 57 | U | N | 46,XX(19)/nonclonal(1) | 3A |
| 10-0090 | M | 58 | M | N | 46,XY(29)/nonclonal(1) | 3A |
| 9-0520 | M | 47 | U | N | 46,XY[20] | 3A, 4A |
| 10-1323 | M | 55 | U | N | 46,XY,inv(1)(p31q32),del(6)(q16q22),del(11)(q22q23)[14]/46,XY[6] | 3A |
| 6-0921 | F | 68 | NA | N | 47,XX,+12(4)/46,XX(8) | 4A |
| 15-1600 | M | 94 | M | T | 47,XY,+12[9]/47,sl,del(10)(q24q26)[11] | 4A |
| 5-0996 | F | 62 | U | N | 47,XX,+X(3)/46,XX(27) | 4A |
| 10-1072 | M | 47 | M | N | 49,XY,+12,+18,+19[2]/49,idem,t(X;13)(p11.4;q14)[15]/46,XY[3] | 4A |
| 16-3489 | M | 76 | U | T | 47,XY,+12[20,two w/nonclonal abnormalities] | 4A |
| 9-1726 | F | 60 | U | T | 46,XX,del(11)(q21q23),inc[cp3]/46,XX[1] | 4A |
| 10-1385 | M | 45 | U | N | 47,XY,+12,t(14;19)(q32.3;q13.2)[4]/46,XY[15]/nonclonal[1] | 4A |

M (mutated IGHV mutational status); U (unmutated IGHV mutational status); T (treated patient); N (treatment-naïve patient); NA (information not available); * (representative immunoblots are shown in Fig. 3B)

### Supplementary Figures

#### Supplementary Figure 1

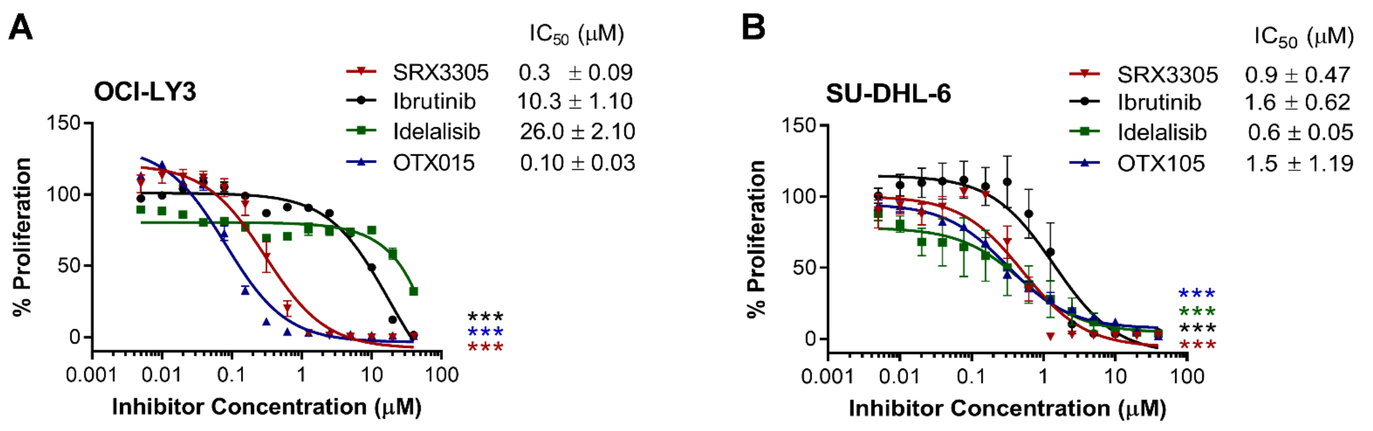


**Figure S1. SRX3305 inhibits DLBCL proliferation.** Two DLBCL cell lines, OCI-LY3 **(A)** and SU-DHL-6 cells **(B),** representative of ABC- and GC-DLBCL subtypes, respectively were treated with increasing doses of SRX3305 (BTK/PI3K/BET inhibitor), ibrutinib (BTK inhibitor), idelalisib (PI3K inhibitor) or OTX015 (BET inhibitor) for 72 h. Proliferation was assessed via MTS assay and results given as % proliferation normalized to vehicle. Error bars indicate SEM (n=3 independent experiments per cell line). Dose-dependent decrease in cell proliferation is observed with asterisks denoting significance (*** $P < 0.001$). $IC_{50}$ values are noted for each inhibitor within the in-figure legends (mean ± SEM).

**Supplementary Figure 2**

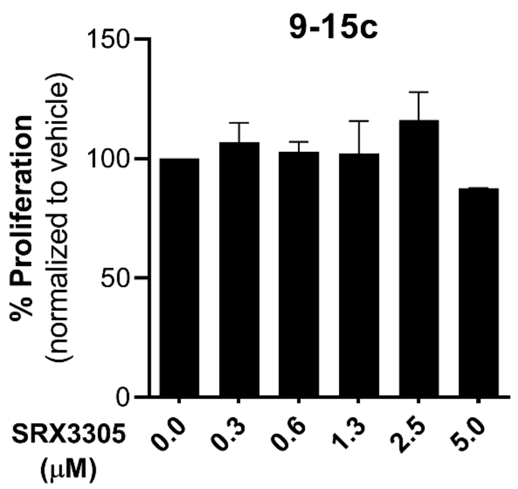


**Figure S2. SRX3305 is not toxic to bystander stromal cells.** Murine 9-15c bone marrow-derived stromal cell lines were treated for 72 h with DMSO vehicle or increasing concentrations of SRX3305 as indicated. MTS assay was done, and stromal cell proliferation is represented relative to vehicle control. Results are given as mean ± SEM (n=3 - 4 independent experiments).

**Supplementary Figure 3**

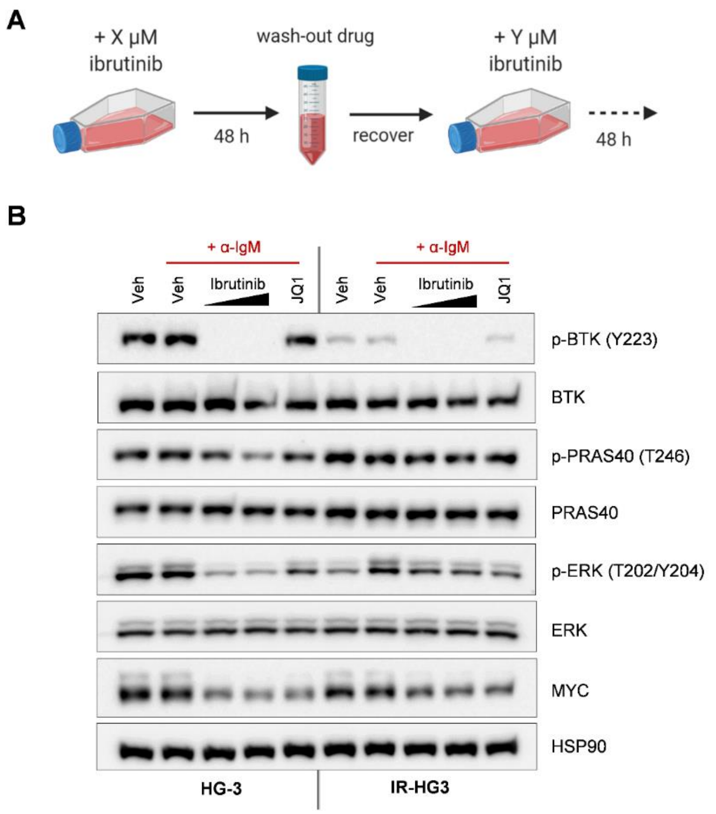


**Figure S3. Generation of an ibrutinib-resistant HG-3 cell line model. (A)** Diagram outlining steps taken to render HG-3 cell resistant to ibrutinib. HG-3 cells (1 $e^6$/mL) were treated with increasing doses of ibrutinib for 48 h at a time, then drug was washed out and the cells were left to recover to ≥ 80% viability before successive dosing (---►). Ibrutinib dose was not increased unless cells successfully recovered within 48 h. Once HG-3 cells remained viable in 25 µM ibrutinib, they were used for further analysis. **(B)** Whole cell lysates of parental (HG-3) and ibrutinib-resistant HG-3 (IR-HG3) cells were collected for immunoblot analysis of the indicated proteins (n=2 independent experiments per cell model). Representative immunoblots are shown.
